# *Wolbachia* effects on Rift Valley fever virus infection in *Culex tarsalis* mosquitoes

**DOI:** 10.1101/135889

**Authors:** Brittany L. Dodson, Elizabeth S. Andrews, Michael J. Turell, Jason L. Rasgon

## Abstract

Innovative tools are needed to alleviate the burden of mosquito-borne diseases, and strategies that target the pathogen instead of the mosquito are being considered. A possible tactic is the use of *Wolbachia*, a maternally inherited, endosymbiotic bacterium that can suppress diverse pathogens when introduced to naive mosquito species. We investigated effects of somatic *Wolbachia* (strain *w*AlbB) infection on Rift Valley fever virus (RVFV) in *Culex tarsalis* mosquitoes. When compared to *Wolbachia*-uninfected mosquitoes, there was no significant effect of *Wolbachia* infection on RVFV infection, dissemination, or transmission frequencies, nor on viral body or saliva titers. Within *Wolbachia*-infected mosquitoes, there was a modest negative correlation between RVFV body titers and *Wolbachia* density, suggesting that *Wolbachia* may suppress RVFV in a density-dependent manner in this mosquito species. These results are contrary to previous work in the same mosquito species, showing *Wolbachia*-induced enhancement of West Nile virus infection rates. Taken together, these results highlight the importance of exploring the breadth of phenotypes induced by *Wolbachia*.

**Author Summary:** An integrated vector management program utilizes several practices, including pesticide application and source reduction, to reduce mosquito populations. However, mosquitoes are developing resistance to some of these methods and new control approaches are needed. A novel technique involves the bacterium *Wolbachia* that lives naturally in many insects. *Wolbachia* can be transferred to uninfected mosquitoes and can block pathogen transmission to humans. Additionally, *Wolbachia* is maternally inherited, allowing it to spread quickly through uninfected field populations of mosquitoes. We studied the impacts of *Wolbachia* on Rift Valley fever virus (RVFV) in the naturally uninfected mosquito, *Culex tarsalis*. *Wolbachia* had no effects on the ability of *Culex tarsalis* to become infected with or transmit RVFV. High densities of *Wolbachia* were associated with no virus infection or low levels of virus, suggesting that *Wolbachia* might suppress RVFV at high densities. These results contrast with our previous study that showed *Wolbachia* enhances West Nile virus infection in *Culex tarsalis.* Together, these studies highlight the importance of studying *Wolbachia* effects on a variety of pathogens so that control methods are not impeded.

## Introduction

Globally, mosquito-borne diseases are a major health burden. To decrease mosquito populations, control programs often use integrated vector management practices including adulticide and larvicide application, source reduction, and biological control [1]. However, these mosquito control methods are losing efficacy due to increasing insecticide resistance and changes in mosquito behavior [2–4]. With these concerns, novel and sustainable control methods are under investigation, including strategies that target the pathogen instead of the mosquito [5,6]. *Wolbachia* is a maternally-inherited endosymbiotic bacterium that infects a large number of insects and other invertebrates [7]. Infection by *Wolbachia* is not innocuous; its presence within a host can cause broad effects on host physiology. For example, natural *Wolbachia* infections in fruit flies protect against pathogen-induced mortality [8,9]. When experimentally transferred to uninfected mosquitoes, *Wolbachia* can suppress infection or transmission of viruses, *Plasmodium* parasites, and filarial nematodes [10–13]. *Wolbachia* also manipulates host reproduction in ways that allow it to spread through and persist in insect populations [14].

Investigations using *Wolbachia*-infected mosquitoes as a control method for dengue virus are underway [15], and field trials in Australia have indicated that *Wolbachia* can spread to near-fixation in naturally uninfected populations of *Aedes aegypti* mosquitoes [16,17]. These *Wolbachia-*infected *Ae. aegypti* populations can persist years after release, and mosquitoes retain the dengue virus-blocking phenotype [18]. Similar field experiments are being conducted in several other countries, but not all have reported successful replacement of the uninfected population with *Wolbachia*-infected mosquitoes [19].

The effects of *Wolbachia*-induced pathogen interference may differ depending on mosquito species, *Wolbachia* strain, pathogen type, and environment conditions [20–22]. For example, in *Anopheles gambiae*, transient somatic infection of the *Wolbachia* strain *w*AlbB inhibits *Plasmodium falciparum* but enhances *Plasmodium berghei* parasites [22,23]. Enhancement phenotypes have been observed in *Anopheles, Culex*, and *Aedes* mosquitoes, and across several malaria species and virus families [20,22,24–27]. Thus, it is important to examine the range of *Wolbachia*-induced phenotypes so that efficacy of disease control efforts using *Wolbachia*-induced pathogen interference are not impeded.

Previous work has demonstrated that transient *Wolbachia* infections in *Culex tarsalis* enhance West Nile virus (WNV) infection rates. To better understand the range of *Wolbachia*-induced phenotypes, we investigated the effects of *Wolbachia* on Rift Valley fever virus (RVFV) infection in *Cx*. *tarsalis*. RVFV is a member of the genus *Phlebovirus* in the family Bunyaviridae and is predominately a disease of domestic ruminants that causes severe economic losses in the livestock industry and human morbidity in Africa and the Middle East [28–30]. Additionally, models and laboratory studies have suggested the United States may have environmental conditions and mosquito vectors that would permit RVFV introduction and invasion [31–34]. *Culex tarsalis* are abundant in the western U.S. and are highly competent laboratory vectors for RVFV [33–35]. We assessed the ability of *Wolbachia* to affect RVFV infection, dissemination, and transmission within *Cx. tarsalis* at two time points and evaluated relationships between viral titer and *Wolbachia* density in mosquitoes.

## Materials and Methods

### Ethics statement

Mosquitoes were maintained on commercially available human blood using a membrane feeder (Biological Specialty Corporation, Colmar, PA). RVFV experiments were performed under biosafety-level 3 (BSL-3) and arthropod-containment level 3 (ACL3) conditions.

Research at the U.S. Army Medical Research Institute of Infectious Diseases (USAMRIID) was conducted under an Institutional Animal Care and Use Committee (IACUC) approved protocol in compliance with the Animal Welfare Act, PHS Policy, and other federal statutes and regulations relating to animals and experiments involving animals. This facility where this research was conducted is accredited by the Association for Assessment and Accreditation of Laboratory Animal Care, International and adheres to the principles stated in the *Guide for the Care and Use of Laboratory Animals*, National Research Council, 2011. The USAMRIID IACUC specifically approved this study.

### Mosquitoes and *Wolbachia*

The *Culex tarsalis* colony used for all experiments was derived from field mosquitoes collected in Yolo County, CA in 2009. Mosquitoes were reared and maintained at 27°C ± 1°C, 12:12 hr light:dark diurnal cycle at 80% relative humidity in 30×30×30 cm cages. The *w*AlbB *Wolbachia* strain was purified from *An. gambiae* Sua5B cells, according to published protocols [36]. *Wolbachia* viability and density was assessed using the LIVE/DEAD BacLight Bacterial Viability Kit (Invitrogen, Carlsbad, CA) and a hemocytometer. The experiment was replicated three times and *w*AlbB concentrations were as follows: replicate one, 2.5 × 10^9^ bacteria/ml; replicate two, 2.5 × 10^9^ bacteria/ml; replicate three, 5.0 × 10^9^ bacteria/ml.

Two- to 4-day-old adult female *Cx. tarsalis* were anesthetized with CO_2_ and intrathoracically injected with approximately 0.1 μl of either suspended *w*AlbB or Schneider’s insect media (Sigma Aldrich, Saint Louis, MO) as a control. Mosquitoes were provided with 10% sucrose *ad libitum* and maintained at 27°C in a growth chamber.

### Vector competence for RVFV

RVFV strain ZH501 was isolated from the blood of a fatal human case in Egypt in 1977 [37]. Adult female Syrian hamsters were inoculated intraperitoneally with 0.2 ml of a suspension containing RVFV in diluent (10% heat-inactivated fetal bovine serum in Medium 199 with Earle’s salts [Invitrogen], sodium bicarbonate, and antibiotics) containing approximately 10^5^ plaque-forming units (PFU) per ml of RVFV. Approximately 28–30 hr post-inoculation, infected hamsters were anesthetized with a suspension of ketamine, acepromazine, and xylazine. A single viremic hamster was placed across two 3.8-liter cardboard cages containing either *Wolbachia*-infected *Cx. tarsalis* or control-injected *Cx. tarsalis,* treatments to which the experimenter was blinded. Mosquitoes were allowed to feed for one hour. After this period, hamsters were removed, a blood sample taken to determine viremia, and hamsters were euthanized.

After feeding, mosquitoes were anesthetized with CO_2_ and examined for feeding status; partially or non-blood fed females were discarded. For all replicates, one blood fed mosquito from each treatment was sampled to test for input viral titers. Mosquitoes were sampled at 7 and 14 days post-blood feeding, where they were anesthetized with CO_2,_ and had their legs removed. Bodies and legs were placed separately into 2-ml microcentrifuge tubes (Eppendorf, Hauppauge, NY) containing 1 ml of mosquito diluent (20% heat-inactivated fetal bovine serum [FBS] in Dulbecco’s phosphate-buffered saline, 50 μg/ml penicillin streptomycin, and 2.5 μg/ml fungizone). Prior to placement into microcentrifuge tubes, saliva was collected from mosquito bodies on day 14 by positioning the proboscis of each mosquito into a capillary tube containing approximately 10 μl of a 1:1 solution of 50% sucrose and FBS. After 30 minutes, the contents were expelled in individual microcentrifuge tubes containing 0.3 ml of mosquito diluent, and bodies were placed in individual microcentrifuge tubes containing 1 ml of mosquito diluent. A 5 mm stainless steel bead (Qiagen, Valencia, CA) was placed into microcentrifuge tubes containing mosquito bodies and legs, homogenized in a mixer mill (Retsch, Haan, Germany) for 30 seconds at 24 cycles per second, and centrifuged for 1 minute at 8000 rpm. All mosquito bodies, legs, and saliva were stored at -80°C until assayed.

Samples were tested for RVFV infectious particles by plaque assay on Vero cells according to previous published protocols [38]. Serial dilutions were prepared for all mosquito body, leg, and saliva samples. One hundred microliters of each dilution was inoculated onto Vero cell culture monolayers. Inoculated plates were incubated at 37°C for 1 hr and an agar overlay was added (1X EBME, 0.75% agarose, 7% FBS, 1% penicillin streptomycin, and 1% nystatin). Plates were incubated at 37°C for 4 days and then a second overlay (1X EBME, 0.75% agarose, and 4% neutral red) was added. Plaques were counted 24 hr after application of the second overlay and titers calculated.

### Quantitative real-time PCR of *Wolbachia* density

To evaluate relationships between *Wolbachia* density and RVFV titer, we measured *w*AlbB levels in individual mosquitoes. DNA was extracted from 200 μl of mosquito body homogenate using the DNeasy blood and tissue kit (Qiagen) and used as template for qPCR on a Rotor-Gene Q (Qiagen) with the PerfeCta SYBR FastMix kit (Quanta Biosciences, Beverly, MA) or on ABI 7500 with Power SYBR green master mix (Applied Biosystems, Foster City, CA). The qPCR assays were performed in 10μl reactions and amplification was carried out using a standardized program at 95°C for 5 min, and 40 cycles of 95°C for 10 sec, 60°C for 15 sec, and 72°C for 10 sec. *Wolbachia* DNA was amplified with primers Alb-GF and Alb-GR [39] and was normalized to the *Cx. tarsalis* actin gene by using qGene software [24,40]. qPCRs were performed in duplicate.

### Statistical analyses

Infection, dissemination, and transmission rates were compared between *Wolbachia*-infected and control *Cx. tarsalis*, and between replicates with Fisher’s exact tests. Due to violations of assumptions needed for parametric tests, Mann-Whitney U was used to compare the following data sets: RVFV body titers between *Wolbachia*-infected and control mosquitoes, RVFV body titers between RVFV-positive saliva and RVFV-negative saliva, RVFV body titers over time, and *Wolbachia* density over time. Unpaired t-tests were used to analyze data that passed normality tests, including the comparison of RVFV saliva titers between *Wolbachia*-infected and control mosquitoes. To determine relationships between *Wolbachia* density and RVFV body titer, the Spearman rank correlation test was used, as assumptions for Pearson correlation were violated. All statistical analyses were performed in GraphPad Prism version 7 for Windows (GraphPad Software, San Diego, CA).

## Results

### Vector competence for RVFV

For all replicates, one blood fed mosquito from each treatment was tested for input RVFV titers on the day of blood feeding. Time 0 results for *Wolbachia*-infected *Cx. tarsalis* were as follows: replicate 1, 2.50 × 10^2^; replicate 2, 7.00 × 10^6^; replicate 3, 1.00 × 10^2^.^0^. Time 0 results for control *Cx. tarsalis* were as follows: replicate 1, 5.00 × 10^2^; replicate 2, 1.05 × 10^7^; replicate 3, 1.00 ×10^2^.^0^. Viremias in the three hamsters were 10^4^, 10^9^, and 10^3^ PFU/ml, respectively.

To determine RVFV vector competence of *Wolbachia*-infected and *Wolbachia*-uninfected *Cx. tarsalis*, we examined frequencies of RVFV-positive bodies, legs, and saliva (Fig. 1). Infection rate is the proportion of mosquito bodies that contained infectious RVFV. Dissemination and transmission rates are the proportion of infected mosquitoes with RVFV positive legs and saliva, respectively. Additionally, transmission rates are also displayed as the proportion of all tested mosquitoes with RVFV positive saliva. Three replicates were performed, and individual data from those experiments are available in Table S1. Hamster viremia in replicate three was low and resulted in low mosquito infection rates. Replicate two infection frequencies were significantly higher than replicate one for both treatments and at both day 7 and day 14 (P < 0.0001). However, across replicates and time points, *Wolbachia*-infected *Cx. tarsalis* infection, dissemination, and transmission rates did not differ significantly from *Wolbachia*-uninfected *Cx. tarsalis* (Fig. 1, Table S1). Thus the data was pooled for further analysis.

**Fig. 1.**
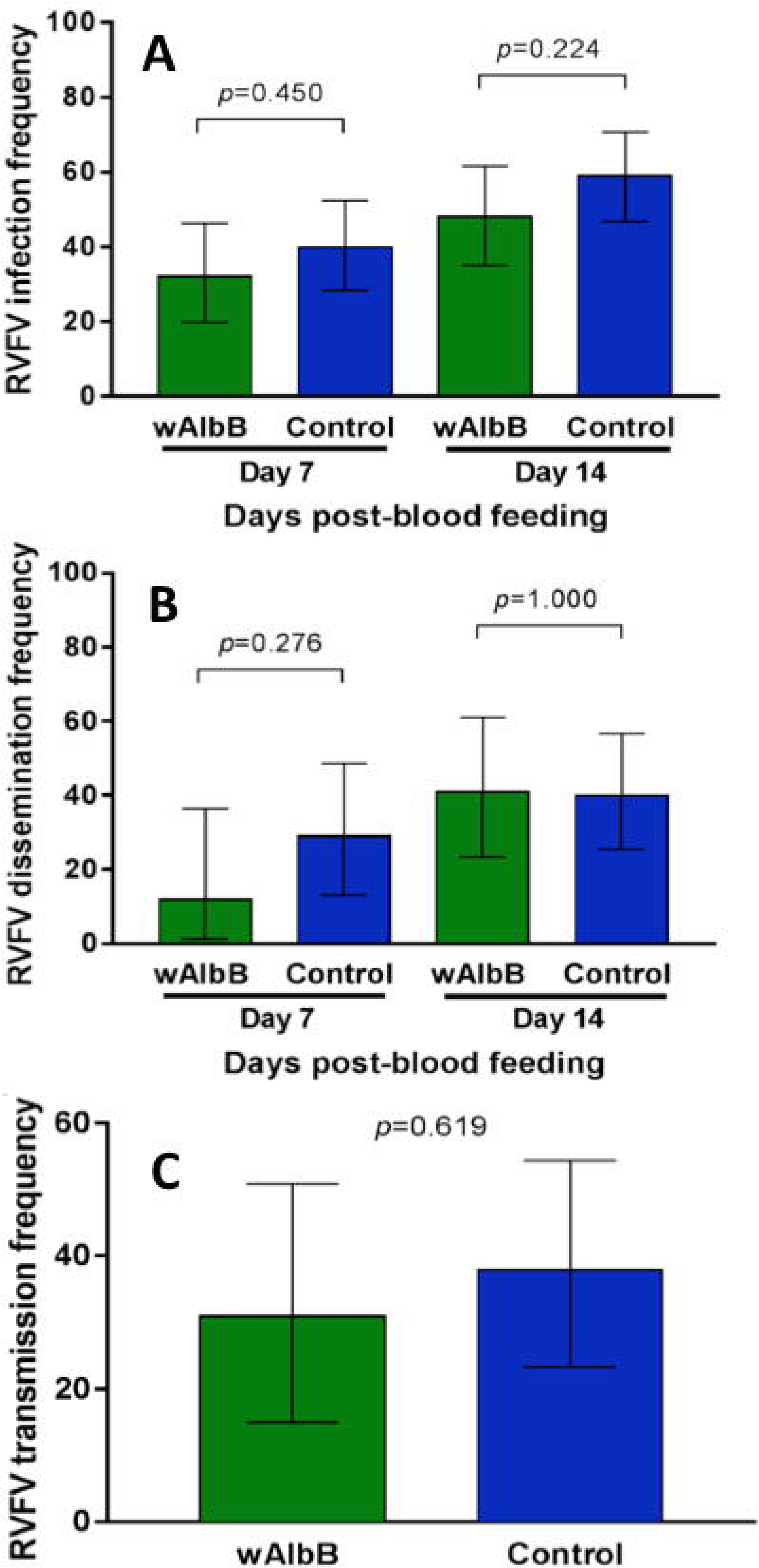
Effects of *Wolbachia* infection on RVFV vector competence frequencies in *Cx. tarsalis*. RVFV infection 7 and 14 days post-feeding (A), dissemination 7 and 14 days post-feeding (B), and transmission rates 14 days post-feeding (C) were compared between *Wolbachia*-infected and control *Cx. tarsalis.* Bars represent data pooled from three replicates. Error bars denote binomial confidence intervals. See Table S1 for replicate-specific analyses.

RVFV body and saliva titers were determined for *Wolbachia*-infected and control *Cx. tarsalis*. There were no significant differences in RVFV body titer or saliva titer between *Wolbachia*-infected and control *Cx. tarsalis* at either day 7 or day 14 (Fig. 2). Additionally, both *Wolbachia*-infected and uninfected *Cx. tarsalis* that transmitted RVFV had significantly higher RVFV body titers than non-transmitting mosquitoes (Fig S1).

**Fig. 2.**
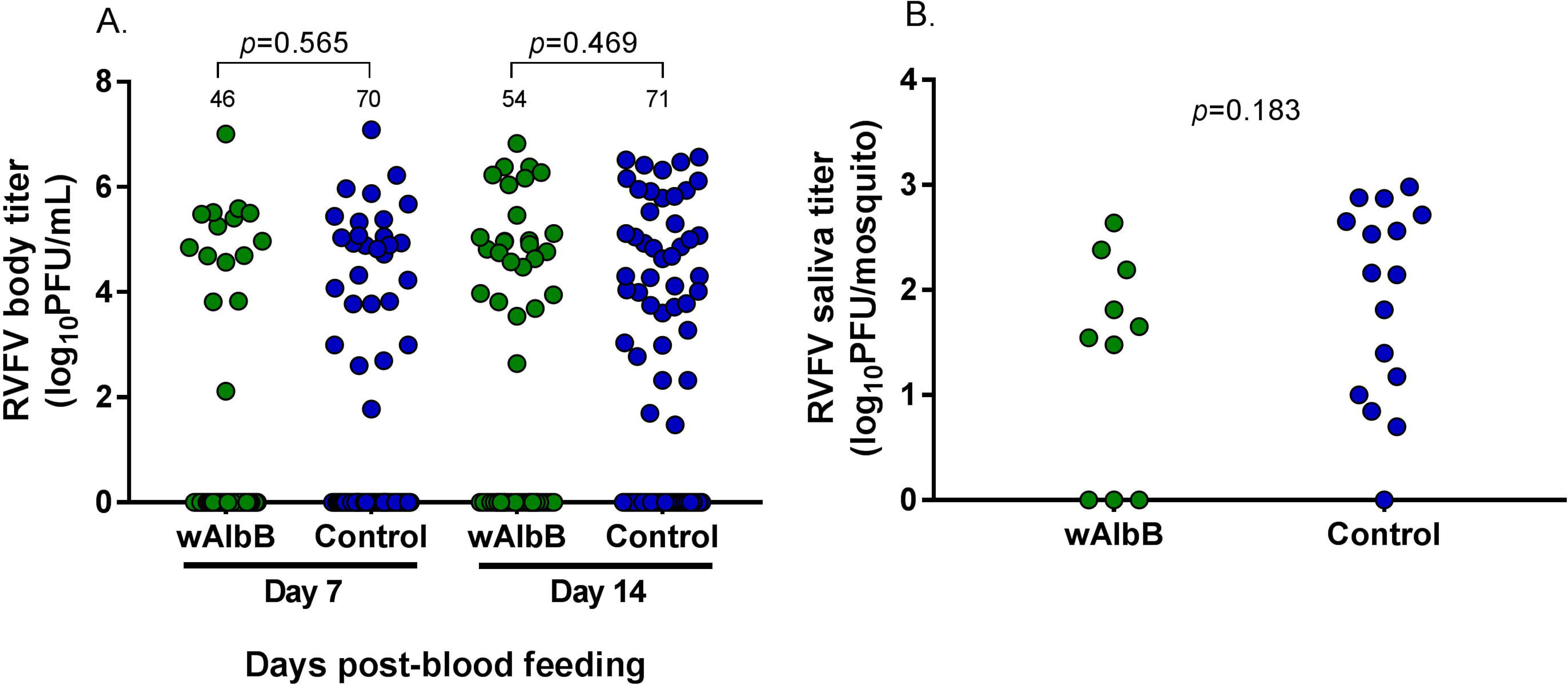
Comparison of RVFV body and saliva titers between *Wolbachia*-infected and control *Cx. tarsalis*. At both 7 and 14 days post-blood meal, there are no significant differences in RVFV body titers of *Wolbachia*-infected *Cx. tarsalis* compared to control *Cx. tarsalis*. All replicates are combined in this figure; separate replicates are provided in supplementary materials (Fig S2). Bars represent medians and bolded numbers above the data points denote sample sizes.

### Quantitative real-time PCR of *Wolbachia* (*w*AlbB) density

*Wolbachia* density in each mosquito was determined by qPCR. We analyzed relationships between *Wolbachia* density and RVFV body titer and combined data from all replicates. Overall, there was a moderate, negative correlation between *Wolbachia* density and RVFV body titer at both day 7 and 14 (Fig. 3). *Wolbachia* density was also compared across time; *Wolbachia* concentration at day 14 was significantly higher than at day 7, consistent with *Wolbachia* replication in mosquitoes (Fig S3).

**Fig. 3.**
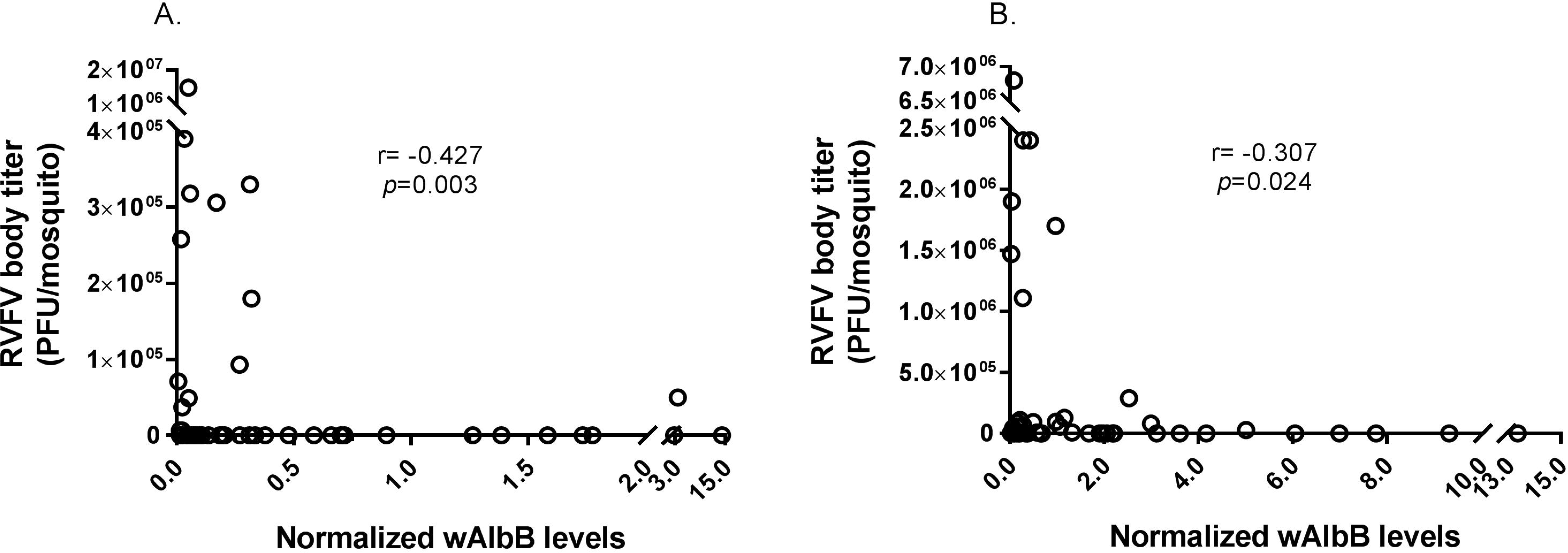
Correlation between RVFV body titer and *Wolbachia* levels in *Cx. tarsalis*. *Wolbachia* levels were normalized to the host gene actin. Normalized *Wolbachia* levels and RVFV body titer for each mosquito were plotted and analyzed with the Spearman rank correlation test to determine relationships. There was a moderate, negative correlation between RVFV body titer and *Wolbachia* levels at both day 7 (A) and day 14 (B) post-blood feeding (Fig. 3). Data for all replicates were combined; see Table S2 for replicate-specific raw data.

## Discussion

*Wolbachia* infection can have varied effects on viruses and parasites transmitted by mosquitoes. These effects can include moderate to complete pathogen inhibition, as well as pathogen enhancement [17,22,24,41,42]. In a previous study, we found that *Wolbachia* strain *w*AlbB enhanced WNV infection frequency in *Cx. tarsalis* [24], although in that study, viral infection titers were not measured. To understand how widespread the *Wolbachia*-induced enhancement phenotype is in *Cx. tarsalis*, we studied *w*AlbB effects on RVFV, an important arthropod-borne virus with potential to invade the United States [43,44]. In contrast to our previous results, we found that *w*AlbB did not affect RVFV body or saliva titers, nor RVFV infection, dissemination, or transmission frequencies in *Cx. tarsalis*.

*Wolbachia*-mediated effects on pathogens may depend on *Wolbachia* density. Several studies have reported that high densities of *Wolbachia* are more likely than low densities to block viruses in *Drosophila* spp. and mosquitoes [45–48]. Similarly, we found a moderate, negative correlation between RVFV body titer and *Wolbachia* density. High *Wolbachia* levels were associated with RVFV negative mosquitoes and very low RVFV body titers. The low numbers of mosquitoes at the high *Wolbachia* densities may explain why we did not see a *Wolbachia* effect on population level vector competence measures. However, our correlation data suggests that in this system, *Wolbachia* may suppress RVFV in a density-dependent manner.

In this *Cx. tarsalis*-*w*AlbB system, we have reported different effects of *Wolbachia* on vector competence for WNV and RVFV [24]. Other studies have found similar differences in *Wolbachia* phenotypes and suggested they may depend on various factors including environmental conditions, and pathogen type [20,49]. RVFV and WNV belong to different virus families and could interact with the mosquito host environment and *Wolbachia* in different ways. For example, a recent study suggested that the mosquito JAK/STAT pathway may not have the same antiviral effects on closely related viruses [50]. Although the mosquitoes in our two studies have the same genetic background, they were reared in separate facilities and may have different microbiomes that may explain differences in vector competence [51]. Another variable that could explain these differences may involve differing blood composition. In the WNV study, we fed mosquitoes on defibrinated bovine blood in a membrane feeder whereas in this study, we fed mosquitoes on live hamsters. Previous studies have suggested that artificial feeding and anticlotting agents may affect various processes within the mosquito [52,53].

Our study was performed with an adult microinjection model that generates mosquitoes transiently infected with *Wolbachia*. It remains to be seen whether this model reflects relationships between *Wolbachia* and viruses in *Cx. tarsalis* in a stable infection system. However, a recent study showed that both stable and transient *w*AlbB infections in *Ae*. *aegypti* produced similar results [45]. This suggests that our transient infection model may correlate with a stable infection in *Cx. tarsalis*.

These studies illustrate the importance of understanding what phenotypes *Wolbachia* influences, and future studies should seek to understand the mechanisms underlying them.

## Acknowledgements

We thank the Cell Culture and Diagnostics Systems Divisions at the United States Army Medical Research Institute of Infectious Diseases (USAMRIID) for providing experimental materials and equipment. We also thank J. Hinson (USAMRIID) for providing technical assistance.

## Figure legends

**S1 Table. Vector competence of *Cx. tarsalis* following a RVFV blood meal.**

RVFV infection, dissemination, and transmission frequencies were compared between *Wolbachia*-infected and control mosquitoes. Replicates are displayed individually.

**S1 Fig. Comparison of RVFV body titers in *Cx. tarsalis* with virus present or absent in the saliva.**

RVFV body titers were compared between mosquitoes that tested positive or negative for RVFV in their saliva. For both *Wolbachia-*infected and control *Cx. tarsalis*, mosquitoes positive for RVFV in the saliva had significantly higher RVFV body titers compared to mosquitoes negative for virus in the saliva There was no significant difference in RVFV body titer of transmitters between *Wolbachia*-infected and control mosquitoes (*p*=0.7692). Data was analyzed with Mann-Whitney U and bars represent medians.

**S2 Fig. Comparison of RVFV body titers between treatments by replicate.**

RVFV body titers were compared between *Wolbachia*-infected and control mosquitoes for replicates 1 (A), 2 (B), and 3 (C). In all replicates, there were no significant differences in RVFV body titer between *Wolbachia*-infected and control mosquitoes. Data did not pass assumptions for normality and were analyzed with Mann-Whitney U, and sample sizes are denoted above data points.

**S3 Fig. *Wolbachia* density over time.**

*Wolbachia* levels for each mosquito, determined by qPCR, were combined across all three replicates. *Wolbachia* levels are significantly higher at day 14 compared to day 7. Due to violations of normality, Mann-Whitney U was used for comparisons, bars are medians, and numbers above data points are sample sizes.

